# DNA methylation sites in early adulthood characterised by pubertal timing and development: A twin study

**DOI:** 10.1101/2023.07.17.549162

**Authors:** Emir Sehovic, Stephanie M. Zellers, Markus K. Youssef, Aino Heikkinen, Jaakko Kaprio, Miina Ollikainen

**Author notes:** Corresponding author – contact.

## Abstract

**Background:** Puberty is a highly heritable and variable trait, with environmental factors having a role in its eventual timing and development. Early and late pubertal onset are both associated with various diseases developing later in life, and epigenetic characterisation of pubertal timing and development could lead to important insights. Blood DNA methylation, reacting to both genotype and environment, has been associated with puberty; however, such studies are relatively scarce. We investigated peripheral blood DNA methylation profiles (using Illumina 450K and EPIC platforms) of 1539 young adult Finnish twins associated with pubertal development scale (PDS) and pubertal age (PA).

**Results:** Fixed effect meta-analysis of the two platforms on 347521 CpGs in common identified 58 CpG sites associated (p < 1 x 10^-5^) with either PDS or PA. All four CpGs associated with PA and 45 CpGs associated with PDS were sex specific. Thirteen CpGs had a high heritability (h2: 0.51-0.98), while one CpG site (mapped to *GET4*) had a high shared environmental component accounting for 68% of the overall variance in methylation at the site. Utilising twin discordance analysis, we found 6 CpG sites (5 associated with PDS and 1 with PA) that had an environmentally driven association with puberty. Furthermore, genes with PDS- or PA-associated CpGs were linked to various developmental processes and diseases, such as breast, prostate and ovarian cancer, while methylation quantitative trait loci (meQTLs) of associated CpG sites were enriched in immune pathways developing during puberty.

**Conclusions:** By identifying puberty-associated DNA methylation sites and examining the effects of sex, environment and genetics, we shed light on the intricate interplay between environment and genetics in the context of puberty. Through our comprehensive analysis, we not only deepen the understanding of the significance of both genetic and environmental factors in the complex processes of puberty and its timing but also gain insights into potential links with disease risks.

## BACKGROUND

Puberty is a necessary developmental phase between childhood and adulthood, enabling sexual reproduction. In addition to pubertal timing and pubertal development differing between the sexes, they are highly variable between individuals of the same sex. Early pubertal development is associated with an increased risk of many diseases in later life, such as breast, endometrial and prostate cancer (1–3). On the other hand, late pubertal development is associated with asthma in both sexes and cervical cancer in females (3). Nevertheless, the mechanisms behind these links remain elusive. Thus, understanding puberty from a genetic and epigenetic perspective may provide further insight into this important period of life and its links with disease risk in later life.

Pubertal timing and development have high heritability, ranging from 37% to 91% (4–7). Studies have identified numerous genetic loci associated with pubertal timing (1,8,9), with a large proportion of the implicated genes involved in neural processes. These identified genetic signals partially explained some diseases, such as breast and prostate cancers, linked to the timing of puberty (1,8,9). Additionally, recent evidence indicates a shift towards an earlier onset of puberty in girls, suggesting that key nongenetic factors play a role in pubertal timing (10,11). Indeed, prenatal growth, diet, maternal smoking, psychological distress in childhood, and exposure to endocrine-disrupting chemicals such as phytoestrogen have been associated with early onset of puberty (12). In addition, although mainly explained through genetic overlap with pubertal timing, high BMI has been associated with early puberty (13,14). Epigenetic marks react to the environment and may provide a mechanism for the environment to act on genome function and ultimately lead to phenotypic consequences. Indeed, DNA methylation, the most studied epigenetic mark, has been associated with pubertal onset (15–17). As the focus of these studies was mainly the change in methylation before and after pubertal timing, there is still a need to identify consistent CpG sites associated with pubertal timing or development in (early) adulthood, where the methylation profile relevant to puberty is more stable than that within early adolescence and could therefore be more reliably linked to disease risk in adulthood. Considering the high heritability of puberty and the wide heritability range of DNA methylation at each CpG site (18), it is necessary to investigate the variance components of DNA methylation sites associated with puberty and to analyse them in the context of methylation quantitative trait loci (meQTLs) (19).

In this epigenome-wide association study (EWAS), we profiled DNA methylation in the blood of young adults to further understand pubertal timing, pubertal development and potential diseases they associate with. We investigated genome-wide DNA methylation and its association with pubertal timing and pubertal development in both sexes. As puberty differs between males and females, we additionally stratified the analyses by sex. We also employed twin study designs to evaluate heritability and within-pair differences and to explore qualitative and quantitative sex differences in DNA methylation at the associated CpG sites. The twin analyses coupled with meQTLs were utilised to gain insight into the genetic and environmental factors affecting methylation values of CpG sites associated with puberty.

## MATERIALS AND METHODS

### Cohorts

This study was based on the participants of two longitudinal and comprehensive birth cohorts of Finnish twins, namely, the FinnTwin12 (twins born in 1983-1987) and FinnTwin16 study (twins born in 1974-1979). Both cohorts have been described in detail elsewhere (20,21). In brief, FinnTwin12 was initiated when the twins were 11-12 years of age, and the follow-up questionnaires were sent at ages 14, 17, and 24, while the FinnTwin16 baseline questionnaire was at the age of 16, with follow-up at the ages of 17, 18, 25, and 35. A subsample of these twins participated in on-site visits, where biological samples were collected, and anthropometric measures were taken (N = 1539, 54.6% female). The current study included both monozygotic (MZ) and dizygotic (DZ) twin pairs who had completed the questionnaires regarding puberty (see below) and had blood DNA methylation data (Illumina 450K or EPIC) in early adulthood available.

### Data on Puberty

The pubertal development scale (PDS) was measured twice in the FinnTwin12 cohort, at ages 12 and 14 years, based on the discrete self-reported markers of puberty: growth spurt, body hair, skin changes, voice change (boys)/breast change (girls) and facial hair (boys)/menarche (girls). PDS was calculated by summing the markers of puberty to obtain a total score, and then the total score was divided by five (22). All individuals with any missing value on PDS variables were excluded from the analysis. Self-reported pubertal age (PA), defined as the age of menarche (girls) or voice break (boys), was obtained from FinnTwin12 participants at the age of 17 and FinnTwin16 participants at the age of 16. All individuals with missing or inconsistent information on PA were excluded. Moreover, individuals who reported that they did not have menarche/voice break yet were coded as the event happening at the age of 17 (13).

### Other Phenotypic Data

For each twin included in the analysis, we also extracted the following items: sex, zygosity, age at questionnaire wave as well as age, smoking status and alcohol consumption at the time of blood sampling. Smoking status was categorised into current, former (abstinent for at least six months) and never smokers, while alcohol consumption was calculated in units of ethanol g/day during the past week, as described previously (23).

### DNA Methylation Data

Blood samples for DNA methylation analyses were collected during in-person visits in adulthood (age range: 21-33.7 and 23.3-42.7 for FinnTwin12 and FinnTwin16, respectively). High molecular weight DNA extracted from whole blood by standard protocols was bisulfite converted with the EZ-96 DNA Methylation-Gold Kit (Zymo) according to the manufacturer’s instructions. Methylation was quantified using Illumina Infinium HumanMethylation450 and EPIC BeadChip platforms that cover more than 450,000 and 850,000 CpG sites, respectively.

The DNA methylation data were preprocessed in the R package *meffil* (24). Bad quality samples were excluded based on the following criteria: i) sex mismatch, ii) median methylation vs. unmethylated signal > 3 standard deviations (SD), iii) failed control probe metrics and if >20% of probes per sample had iv) detection p value > 0.05 and v) bead number < 3. To remove technical variation between the samples, functional normalisation including the control probe principal components was performed, followed by bad quality probe removal: i) detection p value > 0.05, ii) bead number < 3, iii) sex chromosome probes and iv) ambiguously mapped probes (25,26). BMIQ normalisation implemented in the R package *wateRmelon* (27) was then performed to adjust for type 2 probe bias. The same pipeline was used for 450K and EPIC data. The data preprocessing resulted in 390304 and 765385 probes in 450K and EPIC data, respectively. Thirty-nine and 85 samples were excluded after preprocessing in 450K and EPIC data, respectively. Beta values representing DNA methylation were used in all analyses.

### Statistical Analysis

All analyses were performed in R software (version 4.1.1) (28).

#### EWAS Designs

The EWAS analyses were performed using the *limma* package (29). The EWAS on PDS was performed separately on individuals who reported the PDS items at ages 12 and 14. Both sex-stratified and combined models for PDS were performed. The EWAS on PA was performed on the merged cohorts of individuals who reported their PA at age 17 in FinnTwin12 and 16 in FinnTwin16. The EWAS on PA was performed separately in males and females. All EWAS designs are reported in the Supplementary Methods (see Additional file 1). For both 450K and EPIC data, the cell type proportions were calculated using *Flow-Sorted.Blood. EPIC* (30) R package, which is based on a modified version of the Houseman algorithm (31).

We evaluated various combinations of covariates by utilising the AIC to determine the best model fit for the data. This was done using the “selectModel” function from the *limma* package. For each EWAS run, we evaluated the test statistics inflation and bias using the *BACON* package (32). Additionally, for visual investigation of inflation and bias, QQ plots and test-statistic histograms were used, respectively. The QQ and Manhattan plots were created using the *qqman* package (33).

All EWAS models were corrected for smoking, alcohol consumption, age (at the time of blood sampling), cell type proportions, date of BeadChip run and row of the array slide. Additionally, the family ID was included as a random effect to account for the relatedness of twins within a pair. The models that included both sexes were corrected for sex, and the models on PA were corrected for the cohort.

#### EWAS Meta-Analysis

The EWAS results from the 450K and EPIC platforms, based on the same design, were merged using a meta-analysis. A fixed effect meta-analysis with inverse-variance weights was performed using the “rma” function from the *metafor* package (34). The standardised effect sizes and sampling variances were calculated using the “escalc” function. The computed metric was the partial correlation coefficient (35).

The input parameters were the t-statistic, sample size and number of covariates in the regression model. The output parameters were the standardised meta-analysed effect size, p value, Q heterogeneity estimate and its corresponding p value. We corrected the heterogeneity p values using the Benjamini-Hochberg (BH) method. All CpG sites with a corrected Q p value < 0.01 were considered heterogeneous, and a random-effect meta-analysis was performed on such CpG sites.

For the discovery of CpG sites, we used two significance cut-offs on the p values obtained from the meta-analysis: the cut-off recommended for the 450K platform 2.4 × 10^-7^ (36) and a suggestive p value cut-off at 1 x 10^-5^ (37). We annotated all CpG sites with a standardised effect size larger than the absolute value of 0.13 using the *IlluminaHumanMethylation450kanno.ilmn12.hg19* package (38).

#### Analyses of Genetic and Environmental Influences on CpG Sites

To follow up on results from the EWAS, we ran four sets of analyses to further understand the genetic and environmental influences on CpG sites significantly associated with puberty. These analyses, described in detail below, include univariate twin modelling, discordant twin pair analyses, meQTL analysis and analysis of sex differences in CpG sites. Broadly speaking, each analysis provides information on the sources of variation underlying methylation at each CpG site.

#### Univariate Twin Modelling

CpG sites associated with PDS or PA with a p value lower than the suggestive cut-off (p < 1 x 10^-5^) that met the model assumptions and with methylation value SD > 0.05 were assessed by twin modelling (see details in Supplementary Methods - Additional file 1). Univariate twin modelling was performed by the *openMx* R package (39) with age at blood sampling and sex included as covariates to estimate variance components for additive genetic effects (A), common environmental effects (shared exposures and experiences within each twin pair - C), genetic dominance effects (D), and unique environmental effects (nonshared exposures and experiences - E). All potential models (i.e., ACE, ADE, AE, CE, DE, E) were compared to determine the best-fitting set of variance components based on the log-likelihood test.

#### Discordant Twin Pair Analyses

CpG sites associated with PDS or PA with a p value lower than the suggestive cut-off (p < 1 x 10^-5^) were assessed by discordant twin pair analysis. The discordant twin pair design tests whether cotwins who differ in pubertal age or pubertal development also differ in methylation value at each CpG site while naturally controlling for all genetic and environmental confounders shared by the twins within a pair. Discordant twin pair analyses offer the most powerful statistical approach when conducted within MZ twin pairs since the twins in a pair share 100% of their genomes. Here, we additionally pooled MZ and DZ twin pairs together to increase power due to increased sample size compared to the within-pair analyses among MZ pairs only.

We performed a mixed effects model that decomposes the individual-level effect identified in the EWAS into between-pair and within-pair effects. We included a random effect of family and fixed effects of age at blood sampling, sex, and platform as covariates. The between-pair predictor was defined as the average of the exposure within a twin pair. The within-pair predictor (i.e., discordance) was defined as each twin’s value on the exposure subtracted from their cotwin’s value. All pairs were included in the analysis without setting any threshold for within-pair discordance, as they still contribute information to the between-pair effect. Discordant twin pair analyses were conducted in the same subsample for which significant EWAS results were identified.

#### meQTL Analysis

In the context of this study, on the one hand, meQTL SNPs could represent a genetic mechanism that explains the high heritability of a CpG site and, on the other hand, could additionally provide functional information on the SNP-trait association. We used the Genetics of DNA Methylation Consortium (GoDMC) meQTL database (40,41) to explore whether the CpG sites associated with PDS or PA are established meQTL. For each CpG site, we extracted all genome-wide significant (p < 5 x 10^-8^) meQTLs.

#### Sex Differences

Sex differences at the PDS- or PA-associated CpG sites were tested in two ways: differences in methylation values and differences in twin model parameters between the sexes. To test for the differences in DNA methylation value medians between males and females within CpG sites associated with PDS or PA, we performed a Wilcoxon two-sample test using the “wilcox.test” function and corrected the p values using the Bonferroni method. CpG sites with a p value < 0.01 were considered differentially methylated between the sexes. Potential sex differences were also evaluated for all CpGs included in the univariate twin modelling analyses. We explored quantitative and qualitative sex differences, where quantitative sex differences refer to cases where the same genes influence a trait (here, methylation at a particular CpG site) in both sexes but in different magnitudes, and qualitative sex differences refer to cases where different sets of genes influence a trait for each sex (i.e., genes specific to one sex). Importantly, this analysis cannot identify which specific genes influence the trait in each sex, only whether the set of genes as a whole is the same or different between sexes.

An omnibus test was performed to explore whether there were any sex differences in any parameter in the model. The omnibus test compares a model in which every parameter is freely estimated for males and females to a nested model in which all parameter estimates are simultaneously fixed to be equal for both sexes. For all CpG sites with a significant omnibus test, we evaluated each parameter individually to determine which differed between the sexes. The specific sex differences were evaluated by comparing the model in which one parameter was constrained to equality between sexes to a model in which the parameter was freely estimated. P values were generated from likelihood ratio tests and were corrected using the Bonferroni method (function “padjust” in R). The significance cut-off for these adjusted p values was < 0.01 for all tests.

### Pathway Analysis

Ingenuity pathway analysis (IPA) software from QIAGEN Inc. (42) was used to assess whether the PDS- and PA-associated CpG sites and meQTLs were enriched in specific gene ontologies or pathways. The IPA was performed on gene annotated CpG sites with a standardised effect size >|0.13|, using the “Human Methylation 450 v1-2” and the “Ingenuity Knowledge Base” as the reference gene set for CpGs and meQTL SNPs, respectively. The analyses were based only on *Homo sapiens,* and we interpreted the results based on the p value of the associations (significance cut-off was 0.05) with a focus on the canonical pathways and linked diseases or biological functions (“Diseases or function annotation”).

## RESULTS

To identify blood DNA methylation profiles in early adulthood associated with PDS or PA, we performed sex-stratified meta-EWAS on Finnish twins with sample sizes stratified by the array platforms shown in Supplementary Table S1 (see Additional file 3). Furthermore, we also performed EWAS models on PDS using the full sample to determine sex-independent CpG sites. The workflow of the current study is presented in Figure 1. The average age at which the methylation data were obtained was 22.7 years (SD: 1.7) for FinnTwin12 and 28.1 years (SD: 4.3) for the FinnTwin16 cohort. The PDS was obtained from the FinnTwin12 cohort at mean ages of 11.4 (SD: 0.3) and 14.0 (SD: 0.1), while the PA was obtained at mean ages of 16.2 years (SD: 0.1) and 17.6 years (SD: 0.2) from the FinnTwin16 and FinnTwin12 cohorts, respectively. The PDS and PA of the selected individuals (individuals with methylation data included in the study) did not significantly differ from the complete FinnTwin12 and FinnTwin16 cohorts (43). Detailed descriptive statistics on PDS, PA and other relevant variables can be seen in Table 1. A total of 347521 CpG sites were included in the EWAS meta-analyses, with no evidence of inflation or bias in any of the EWAS models (across both platforms, the bias test statistic ranged from -0.213 to 0.142, while the inflation test statistic ranged from 0.9 to 0.995). Overall low heterogeneity was observed between the two platforms (< 0.07% heterogeneous CpGs across all models).

**Figure 1.**
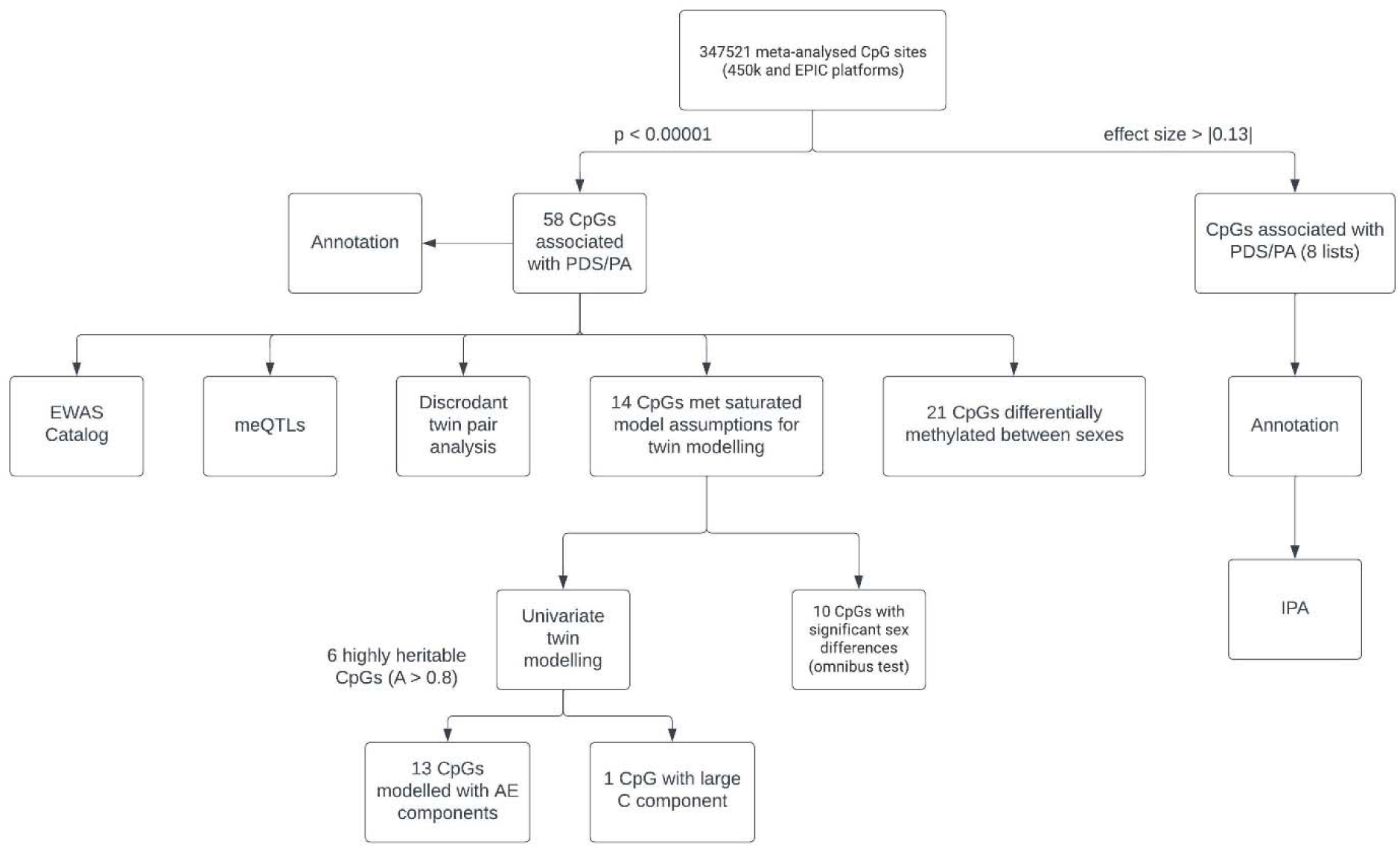
Workflow diagram of the study. Through a meta-analysis of 450K and EPIC platforms on 347521 CpGs, we identified CpG sites associated with PA or PDS at age 12 or 14 (p < 0.00001). We performed univariate twin modelling to assess heritability and to define the proportion of variance attributed to unique and shared environments and investigated potential sex differences in the significant CpG sites. We identified enriched pathways and linked diseases among CpG sites with an absolute effect size > 0.13 by IPA.

**Table 1.**
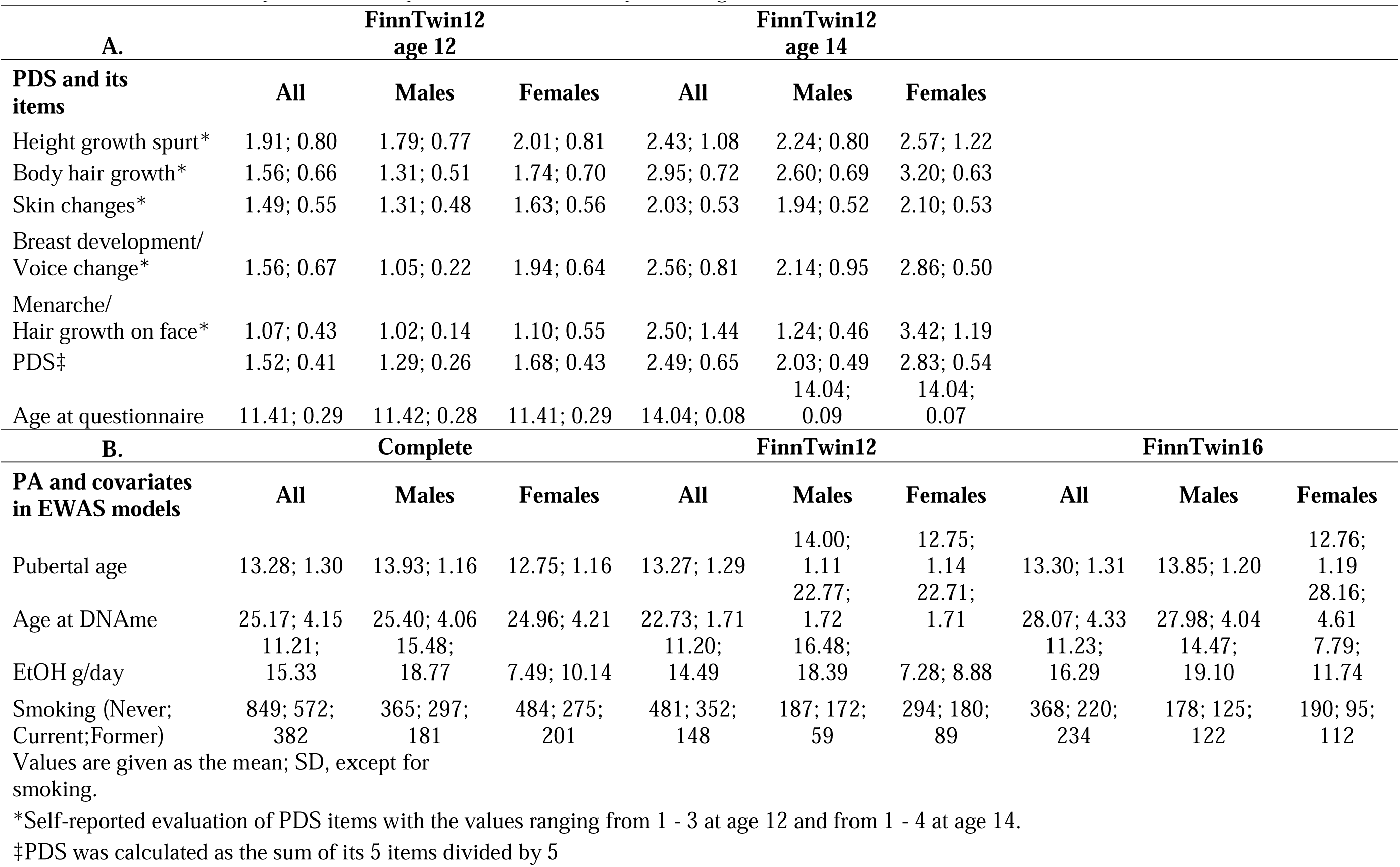
Characteristics of pubertal development scale (PDS) and pubertal age (PA), and covariates in the EWAS.

### Puberty-associated CpG Sites

At the age of 12, PDS was associated with 9 CpG sites (p < 1 x 10^-5^) in both sexes combined, while 13 and 10 were exclusively associated with PDS in males and females, respectively (Table 2, Supplementary Figure S1 and S2 - Additional file 2). Two CpG sites in females (cg07581365, cg06988547) had a lower p value than the threshold of 2.47 x 10^-7^. At the age of 14, no CpG sites were associated with PDS in the model including both sexes (Supplementary Figure S1 and S2 - Additional file 2). However, 12 and 10 CpG sites were associated with PDS in males and females, respectively (Table 2 and Figure 2). One CpG site in males (cg04239863) and one in females (cg20599748) were significantly associated with PDS (p < 2.47 x 10^-7^). A total of 12 of the 54 CpG sites associated with PDS showed significant heterogeneity between the 450K and EPIC platforms (Table 2), indicating lower reliability of the observed associations for those CpG sites. After performing a random effect meta-analysis for these CpG sites, none remained significantly associated with PDS.

**Figure 2.**
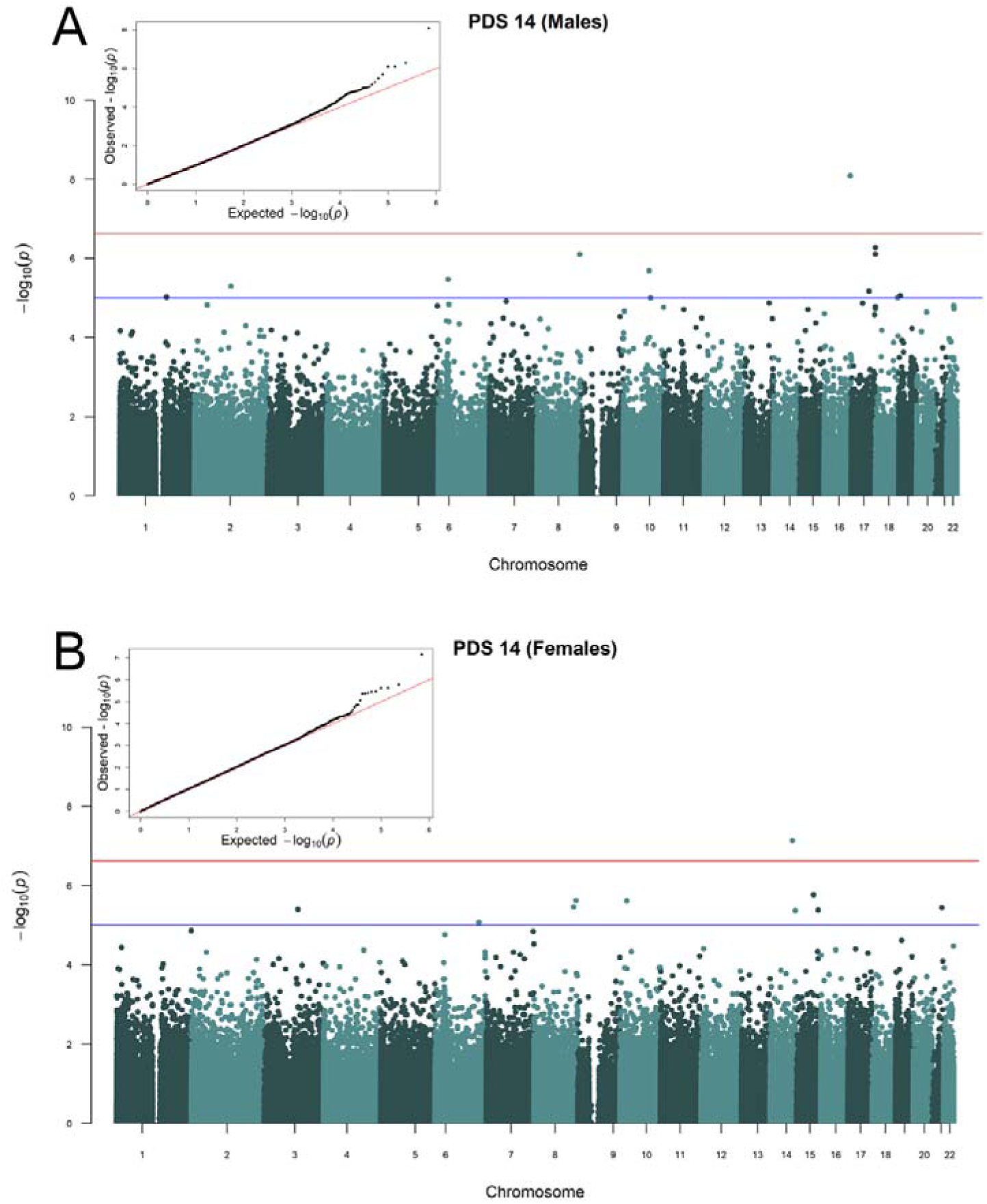
Manhattan plots of the meta-analyses. The Manhattan plots show significant associations between CpG methylation and PDS at age 14 in A) males and B) females. The red line indicates a significance cut-off of 2.4 x 10^-7^ as recommended for the 450K platform, while the blue line indicates a suggestive significance cut-off at 1.0 x 10^-5^. Embedded (top left corner of Manhattan plots) are the QQ plots of meta-analysis p values.

**Table 2.**
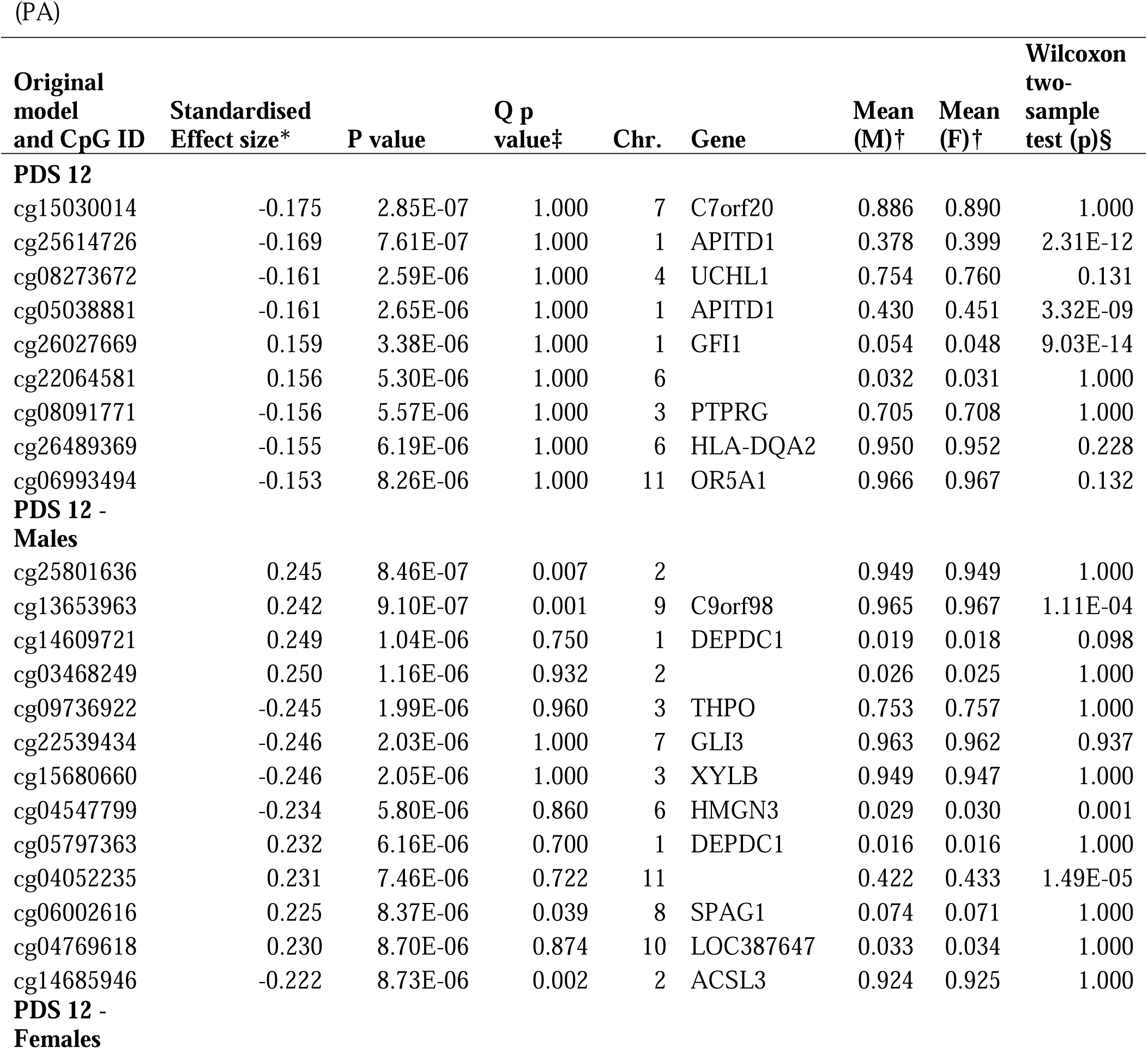
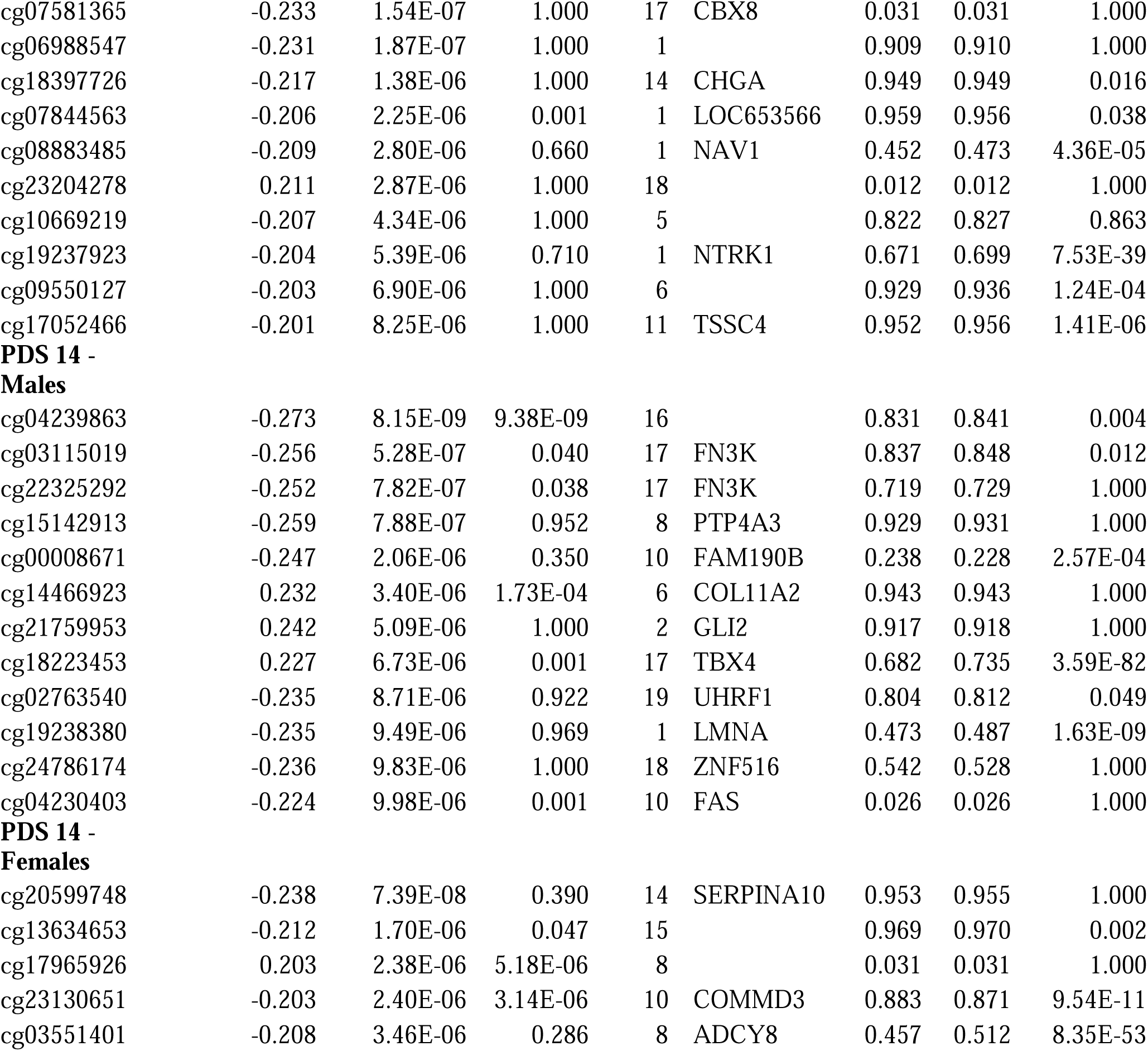
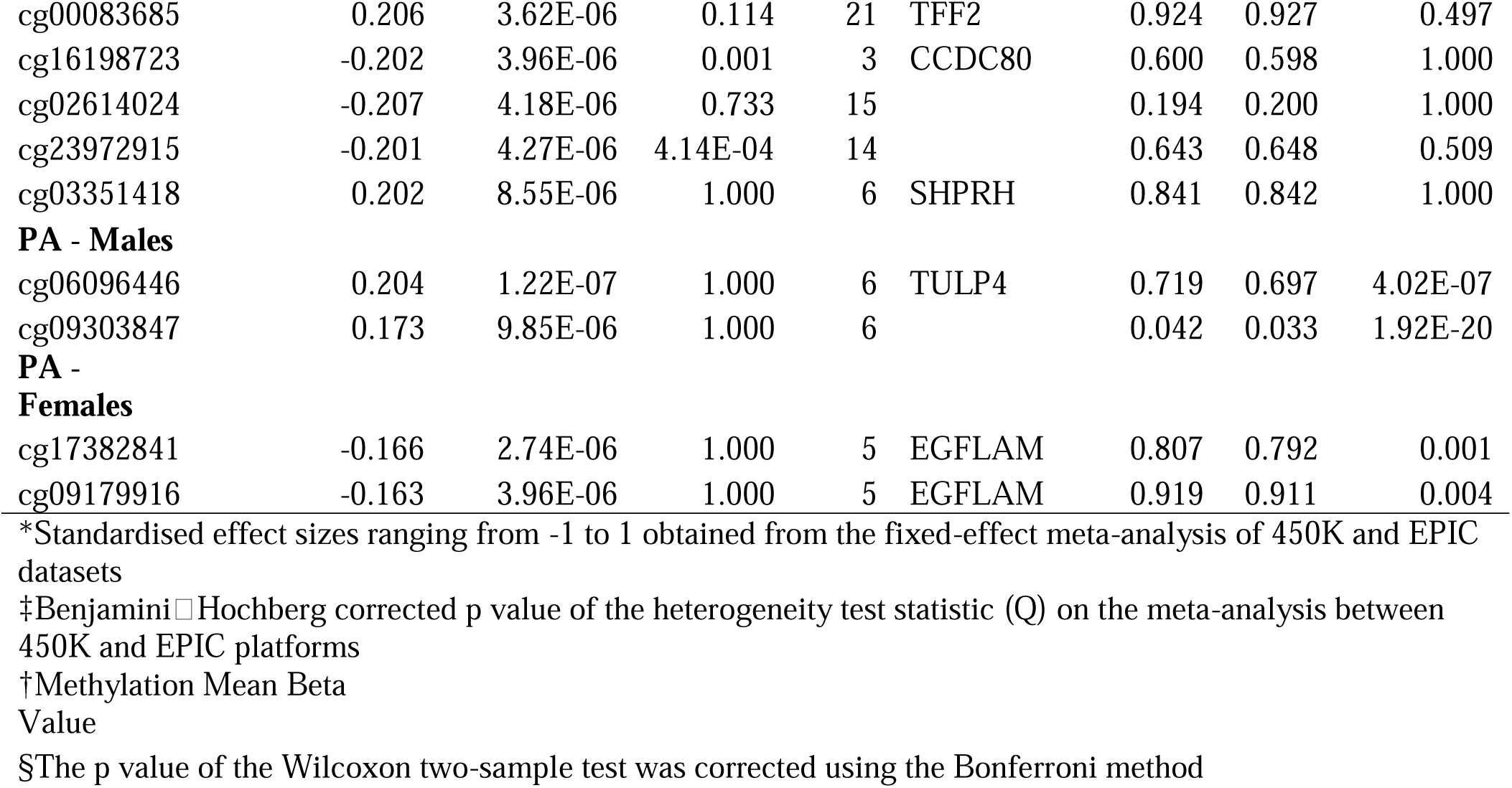
CpG sites significantly associated (p<1x10^-5^) with pubertal development scale (PDS) and pubertal age.

Four CpG sites were associated with PA with p < 1 x 10^-5^: two unique CpGs in males and two in females (Table 2, Supplementary Figure S1 and S2 - Additional file 2). One of the CpG sites associated with PA in males (cg06096446) had a p < 2.47 x 10^-7^. None of the CpGs associated with PA were heterogeneous between the two platforms.

### Genetic and environmental effects underlying the associations

First, as puberty itself as well as DNA methylation are highly heritable, we wanted to investigate the heritability and shared/nonshared environmental components explaining the variance of the 58 CpG sites associated with PDS or PA in early adulthood. We identified 14 CpGs for which both assumptions of the twin models were met, and the CpG had methylation SD > 0.05 (Supplementary Table S2 - Additional file 3). With respect to the univariate models, 13 out of 14 CpG sites were best modelled with only additive genetic (A) and unique environmental components (E) (Table 3). Heritabilities for the AE CpG methylation values ranged from 0.51-0.98, indicating strong additive genetic influences on these CpGs (average heritability was 0.79). Only one of the 14 CpGs (cg15030014, associated with PDS at age 12 in both sexes) also required the inclusion of a shared environmental component (C), indicating strong familial resemblance on this CpG due to nongenetic sources in addition to additive genetic influences (A=0.24, C=0.68).

**Table 3.** Univariate and sex differences twin modelling of 14 CpG sites meeting the twin modelling assumptions.

| CpG Name | MZ Twin Correlation | DZ Twin Correlation | Mean | SD | Best Model | Heritability [95% CI] | Shared Environment [95% CI] | Unique Environment [95% CI] | Sex Differences Identified |
| --- | --- | --- | --- | --- | --- | --- | --- | --- | --- |
| cg02614024 | 0.69 | 0.29 | 0.19 | 0.06 | AE | 0.67 [0.61, 0.72] | - | 0.33 [0.28, 0.39] | Unstandardised genetic and environmental variance components |
| cg03115019 | 0.96 | 0.50 | 0.85 | 0.11 | AE | 0.96 [0.95, NA] | - | 0.04 [NA, 0.05] | None |
| cg03551401 | 0.55 | 0.33 | 0.49 | 0.07 | AE | 0.51 [0.42, 0.58] | - | 0.49 [0.42, 0.58] | Mean structure |
| cg05038881 | 0.83 | 0.44 | 0.44 | 0.06 | AE | 0.84 [NA, NA] | - | 0.16 [0.13, NA] | Mean structure |
| cg06096446 | 0.89 | 0.48 | 0.70 | 0.09 | AE | 0.88 [0.86, 0.90] | - | 0.12 [0.10, 0.14] | Mean structure |
| cg08883485 | 0.94 | 0.52 | 0.46 | 0.08 | AE | 0.94 [0.92, NA] | - | 0.06 [NA, 0.08] | Mean structure |
| cg09179916 | 0.61 | 0.18 | 0.92 | 0.06 | AE | 0.60 [0.53, 0.67] | - | 0.40 [0.33, 0.47] | Mean Structure,<br>unstandardised<br>genetic and<br>environmental<br>variance<br>components |
| cg15030014 | 0.89 | 0.78 | 0.89 | 0.06 | ACE | 0.24 [0.17, 0.33] | 0.68 [NA, 0.75] | 0.08 [0.07, 0.10] | Unstandardised<br>genetic variance<br>component<br>and total<br>phenotypic<br>variance |
| cg16198723 | 0.79 | 0.36 | 0.60 | 0.05 | AE | 0.78 [0.73, NA] | - | 0.22 [NA, 0.27] | None |
| cg17382841 | 0.61 | 0.28 | 0.80 | 0.07 | AE | 0.64 [0.56, 0.70] | - | 0.36 [0.30, 0.44] | Mean structure |
| cg18223453 | 0.81 | 0.48 | 0.71 | 0.05 | AE | 0.77 [0.72, 0.81] | - | 0.23 [0.19, 0.28] | Mean structure |
| cg22325292 | 0.98 | 0.53 | 0.73 | 0.19 | AE | 0.98 [0.97, 0.99] | - | 0.02 [NA, 0.03] | None |
| cg24786174 | 0.97 | 0.55 | 0.53 | 0.14 | AE | 0.97 [0.96, NA] | - | 0.03 [NA, 0.04] | None |
| cg25614726 | 0.77 | 0.42 | 0.39 | 0.05 | AE | 0.78 [0.74, NA] | - | 0.22 [0.19, 0.26] | Mean structure |

Best twin model, standardised variance components and potential identified sex differences are shown

Second, as only 14/58 CpG sites met the twin modelling criteria, we applied a discordant twin pair study design to identify PDS- and PA-associated CpG sites with their methylation likely driven by nongenetic factors. Mixed effects models were run to decompose the effect identified in the EWAS into within-pair and between-pair associations. Significant within-pair associations are consistent with a nongenetic effect. Here, CpGs that were considered differentially methylated within the pairs had a significant within-pair difference in the pooled analysis (MZ and DZ) as well as a suggestive difference in the same direction (p < 0.10) in the MZ-only analyses. Full results are available in Supplementary Table S3 in Additional file 3. We identified six differentially methylated CpGs, five of which were associated with PDS-12 and one with PA. For three CpGs, the MZ twin with a higher PDS-12 score also had higher methylation than their cotwin (cg14609721, cg03468249, cg05797363). For two CpG sites, the MZ twin with a higher PDS-12 score had lower methylation than their cotwin (cg08091771, cg10669219). Finally, for one CpG, the MZ twin with older PA also had higher methylation than their cotwin (cg06096446). These effects suggest an environmentally driven relationship between early pubertal development and DNA methylation. This effect is consistent with a causal relationship but cannot determine the direction of effect (i.e., whether methylation causes earlier puberty or if early puberty causes methylation).

Third, to gain additional insight into the genetic basis of DNA methylation at the 58 CpG sites associated with PDS and PA, we looked for *cis* and *trans* meQTLs from the GoDMC database affecting methylation at these sites. Thirty-nine of the 58 CpG sites had *cis*-meQTLs, and only one CpG had *trans*-meQTLs (Figure 3). The number of associated meQTLs ranged from 1 to 1676 at these 39 CpG sites. Additionally, we found 3 CpG pairs whose methylation was affected by a subset of common SNPs (Figure 3).

**Figure 3.**
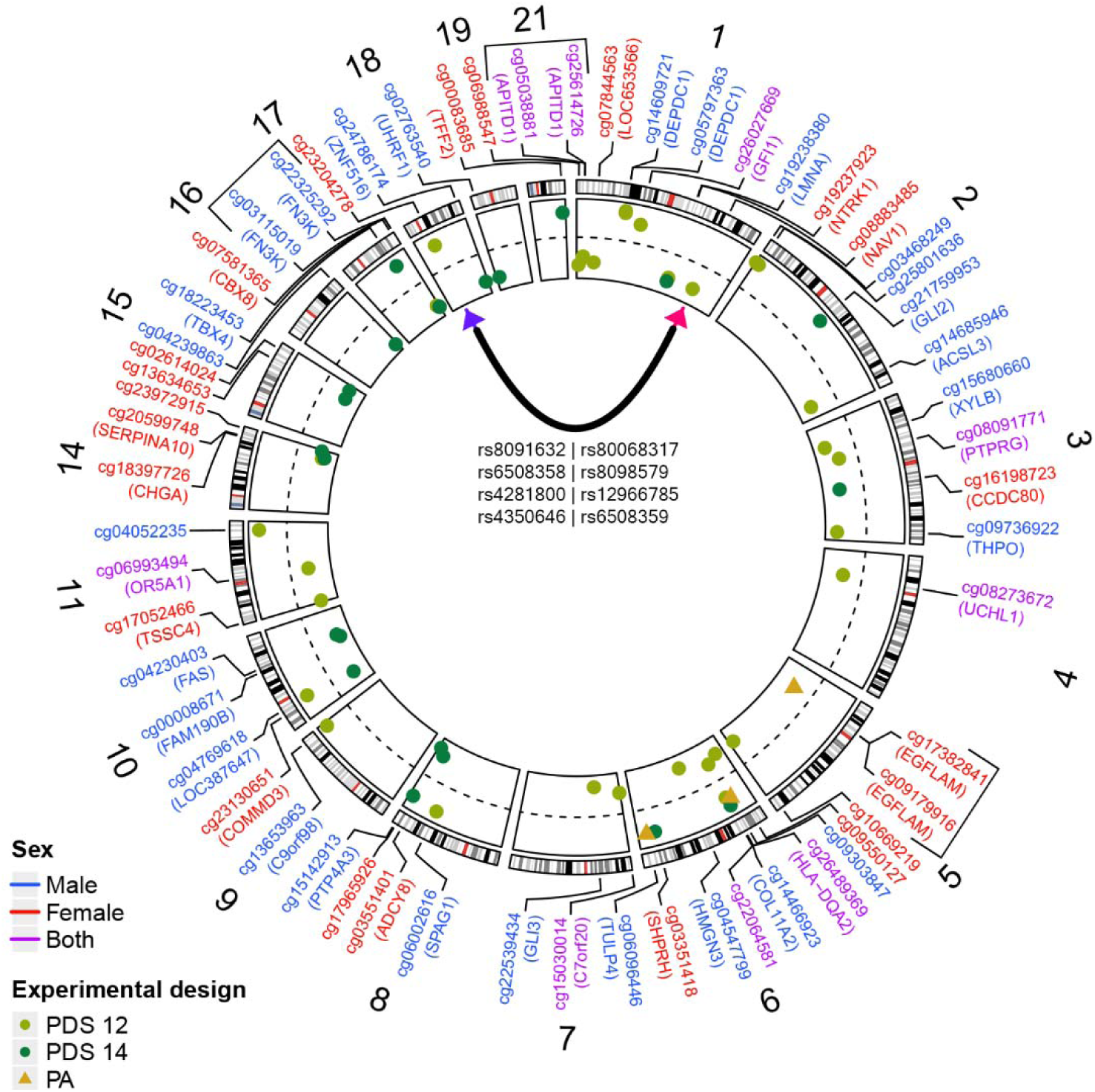
Circos plot showing the chromosomal locations of the 58 CpGs associated with PDS or PA. The inner circle shows the standardised effect sizes of the association (scale ranging from -0.3 to 0.3) with the dashed line representing 0. The shape of the points plotted on the effect size scale determines the variable analysed (PDS or PA) in the model in which the CpG was found, while the colour of the CpG site identifier shows whether the model was performed on males, females or both sexes. The 8 trans-meQTLs of CpG cg08883485 are depicted with the black line inside the circos plot (pink arrowhead pointing to the CpG site and purple arrowhead pointing to the genomic region of the 8 trans-meQTLs). The CpGs with common cis-meQTLs are shown with black square brackets.

### Sex Differences in CpG sites

As we observed overall stronger associations in the sex-stratified models compared to models including both sexes and as there were no common significant CpG sites between the EWAS models in males vs females, we aimed to characterise potential sex differences and similarities in the identified CpG sites. Therefore, we performed a Wilcoxon two-sample test on CpG sites associated with PDS or PA and investigated sex differences by exploiting the properties of opposite-sex dizygotic twin pairs. Based on all included samples in this study, 21 CpG sites were differentially methylated between males and females (Table 2). Three differentially methylated CpGs were obtained from models that included both sexes (associated with PDS at the age of 12).

To investigate sex differences utilising the twin modelling method, we first examined qualitative sex differences (i.e., whether different sets of genes impact CpG methylation in males and females). We also performed an omnibus model on all 14 CpGs included in the twin analyses to determine whether there are any quantitative sex differences (i.e., whether parameter estimates differ between males and females). Qualitative sex differences existed for none of the 14 CpGs, indicating that the same additive genetic influences impact methylation in both males and females. We obtained ten CpG sites (9 based on AE models and 1 ACE model) with quantitative sex differences (Table 3, Supplementary Table S4 - Additional file 3). Three CpGs exhibited significant differences in variance compositions. For both cg09179916 and cg02614024 (associated with PA in females and PDS at 14 in females, respectively), the additive genetic component was larger in females, and the nonshared environmental component was larger in males, but total variance did not differ between the sexes. For cg15030014 (associated with PDS at age 12 in both sexes), both the additive genetic component and the total phenotypic variance were larger in males. In general, we identified sex differences in both methylation and its underlying variation in CpGs associated with puberty in one or both sexes.

### Enrichment Analysis

We obtained relevant canonical pathways and associated diseases/functions using IPA software on PDS- and PA-associated CpG sites with an effect size larger than the absolute value of 0.13 (see Supplementary Table S5 and S6 for the CpG sites - Additional file 3). CpGs associated with PDS at the age of 14 in males were enriched in sperm motility, axonal guidance signalling and synaptogenesis signalling canonical pathways, while in females, the pathway results were less conclusive. Full results of the canonical pathways from IPA for all models can be seen in Supplementary Table S7 (see Additional file 3).

CpGs associated with PA in males were enriched in adipogenesis, choline degradation and thyroid hormone biosynthesis canonical pathways, while in females, they were enriched in tyrosine degradation and choline biosynthesis. Across all sex-specific models, associated CpGs were linked to physiological system development, such as growth of muscle tissue or development of neural cells in PDS 14 models or connective tissue development and tissue morphology in PA (Supplementary Table S8 - Additional file 3). Breast, prostate and ovarian cancers were universally found to be linked to CpGs in the sex-stratified models on PDS both at ages 12 and 14. Intestinal and thyroid cancers were highly significantly linked diseases in CpGs associated with PDS at age 14 in males. In females, the highly significant diseases were cancer of the head (umbrella term) and gastric cancer (Supplementary Table S8 - Additional file 3).

To explore whether the SNPs underlying the methylation associations with puberty are enriched in any relevant processes and functions, we performed IPA on meQTLs of all 39 CpG sites. IPA was performed on 13787 unique meQTLs, and they were enriched in several immune pathways, such as antigen presentation and the T-helper 1 and T-helper 2 pathways, usually driven by genes linked to the human leukocyte antigen (HLA) complex (e.g., *HLA-DQA2* and *HLA-DPA1*). The multiple sclerosis signalling pathway was one of the highly enriched canonical pathways among the meQTLs (Supplementary Table S9 - Additional file 3). Concerning the diseases and functions category of IPA, several diseases were highlighted, such as insulin-dependent diabetes mellitus, rheumatoid arthritis, and endocrine, thyroid and breast cancers.

## DISCUSSION

In a sizable cohort of Finnish twins, we identified methylation values at 58 CpG sites to be positively or negatively associated with PDS or PA when assessed among individuals in early adulthood. We observed both sex-specific associations and associations common to both sexes, with overall stronger associations in sex-stratified models. On average, we identified stronger and more numerous associations with PDS than PA. Moreover, relatively few CpGs overlapped between any of the models.

We considered PDS at age 14 of particular importance; compared to the models at age 12, it is generally closer to pubertal development peak (44), while PA was reported retrospectively, making it more prone to reporting errors. Additionally, CpG sites with methylation associated with PDS at age 14 were frequently located in development-related genes. For instance, male PDS-related hypomethylated cg19238380 is located in the *LMNA* gene, which plays a critical role in the normal development of the peripheral nervous system and skeletal muscle (45,46). Additionally, the CpG site cg03551401, where methylation is negatively associated with PDS in females, is located on the *ADCY8* gene, which contributes to several brain functions, including learning, memory, and drug addiction (47).

Three CpGs, with an effect size larger than the absolute value of 0.13 but below the specified p value thresholds, were also found in two studies that investigated genome-wide DNA methylation profiles in peripheral blood before and after puberty (16,17). The three CpG sites are cg01794929, cg21361322 and cg15028548 (the former two mapped to the *SELM* gene and the latter to *ABI3BP*) and were associated in our study with PDS at 14 in females, PDS at 12 in females and PDS at 14 in males, respectively. Notably, the main difference between our study and the two mentioned studies is that we explored the methylation profiles of young adults explained by pubertal timing and development, while they investigated methylation profiles before and after puberty. This difference is reflected in the observed opposite directionality of effect sizes for the 3 CpG sites.

Overall, canonical pathway enrichment results when both sexes were considered together were less conclusive than sex-specific pathway analyses, in which multiple CpG sites were enriched in endocrine or neuronal development-related pathways. Additionally, an antagonistic enrichment related to choline and tyrosine metabolism was found between males and females among the CpG sites associated with PA. Furthermore, breast, ovarian and prostate cancers were linked to CpG sites across all models, which is consistent with studies showing associations between early puberty and these diseases (1,9). Previously reported diseases related to early puberty, such as endometrial cancer (1,9) or cardiovascular diseases (3), were also linked to CpG sites in some of the models in the current study, such as in the sex-stratified PDS at age 14.

In addition to characterising the genes relevant to the identified CpG sites, we also examined their underlying sources of variation via twin modelling, as both puberty (4–7) and DNA methylation are known to be heritable traits (18,19). The CpG sites reported in this study, which were predominantly best modelled with additive genetic and unique environmental sources of variation, were mostly found to be highly heritable. Generally, methylation values of these CpGs were more heritable than pubertal development or PA themselves, with heritability ranging between 0.51-0.98 for AE CpGs, while for age at menarche or age at voice change, it ranges from 0.50-0.59 (4,5,48). Methylation at only one CpG site (cg15030014), located in the *GET4* gene, showed strong influence from the environment. Interestingly, *GET4* has an enhancer with a SNP (rs9690350) mapped to the gene region of *PDGFA,* which is associated with pubertal onset (49). The same CpG, as well as 47 other significant CpGs, are associated with age (50), which supports our findings on associations with pubertal timing. Additionally, they could be considered markers for epigenetic age acceleration in this critical period of life. It is unclear to what extent the high heritability of CpGs observed here may reflect genetic mechanisms that influence pubertal development, given that the blood sampling occurred after the completion of puberty.

To assess in more detail the potential genetic effects on puberty-associated CpG sites, we performed meQTL analyses. A total of 4507 unique SNPs associated with age of menarche in two large-scale genome-wide association studies (GWAS) (1,9) were identified as meQTLs in our study. They could provide a putative mechanistic explanation for PDS- or PA-associated SNPs identified in the GWA studies, as DNA methylation may impart a mechanistic link for these SNPs and explain their association with age at menarche. A nonzero proportion of SNPs reported by the two studies was found in all our designs, barring PDS at age 14 in males and females combined. The percentage of meQTLs associated with CpGs from the eight models, also found among the 4507 SNPs, ranged from 21.1% to 43.2% (26-1289 meQTLs). The meQTLs of highly heritable CpGs also had a substantial proportion of common meQTLs with GWAS on puberty, ranging from 1% to 36.3% (16-465 meQTLs). Hence, these meQTLs could, in part, reflect genetic mechanisms that influence pubertal development.

Pathway-level analyses on the meQTLs of CpG sites associated with puberty revealed an enrichment in HLA complex-related immune pathways such as antigen presentation and T-helper pathways. A positive association between HLA complex heterozygosity and later pubertal development was reported and was hypothesised to be an evolutionary trade-off between immunocompetence reflected in pathogen resistance and sexual maturation (51). Moreover, an increase in antigen presentation and T-helper responses has been reported as part of immune changes during pubertal development (52). Thus, pathway analyses on meQTLs underlying the methylation associations with puberty demonstrate their involvement in immune processes fully developed during puberty, which might be relevant in terms of susceptibility to various diseases later in life. Furthermore, the IPA of meQTLs revealed some of the same linked diseases as our IPA of differentially methylated CpG sites, such as endocrine, breast and thyroid cancers. Interestingly, the enriched immune pathways were also associated with the central nervous system and could be linked to susceptibility to multiple sclerosis (52), which was strongly enriched in the meQTL IPA analysis.

To rule out the effects of shared genotypes and environment on the observed associations, we performed within-pair analyses. We observed 6 CpG sites with their methylation differing within the pairs, indicating a causal relationship between PDS or PA and methylation at these CpG sites. Two of these CpGs were positively related to PDS, and three were negatively related to PDS, while only one CpG site (cg06096446) with high heritability, whose mapped gene (*TULP4*) is associated with height (53), was positively related to PA. The same CpG site has also been associated with alcohol consumption and disorders in pregnancy (54,55). One of the negatively associated CpGs with a significant within-pair methylation difference (cg08091771) has been associated with maternal BMI and nitrogen dioxide exposure (56,57) and is mapped on *PTPRG,* which is associated with precocious puberty (58). These effects are consistent with a causal environmentally mediated relationship between earlier pubertal development and methylation, although our analyses cannot conclusively determine the direction of causality (i.e., whether methylation causes earlier puberty or whether earlier puberty causes methylation). The rest of the puberty-associated CpG sites did not differ within the pairs, suggesting that the relationship between PDS/PA and CpG methylation is due to genetic or environmental confounding.

Twin modelling results on quantitative sex differences and differential methylation results were concordant: all CpGs identified as having sex differences in the mean structure were also differentially methylated between the sexes. Furthermore, for CpGs with sex differences in variance, the sex with a larger variance from additive genetic sources was also the sex for which the EWAS association was identified. For example, cg09179916 was associated with PA in females, and females had a larger additive genetic component. This convergence of results could suggest a mechanistic role, but further analyses are necessary to determine if methylation at these CpG sites has a functional role related to puberty. In addition to the mechanistic role, the observed sex differences emphasise the importance of sex-stratified models for epigenetic studies of puberty.

We found an overall high heritability across the PDS- and PA-associated CpG sites, which for 6 CpGs was likely due to direct SNP effects observed as meQTLs. However, the methylation discordance within twin pairs revealed that, for some CpG sites, despite their high heritability, environmental factors play an essential role in shaping their methylation, possibly indicating the presence of gene-environment interactions. Finally, the observed sex differences indicate that the same genetic mechanisms, but with different magnitudes in males and females, affect the heritability of CpG sites associated with puberty, giving additional importance to investigating meQTLs in this context.

Our study has both strengths and limitations. The key strengths of our study are the relatively large sample size as well as the inclusion of twin pairs with longitudinal phenotype data on which we were able to separate genetic effects from environmental effects on methylation at the puberty-associated CpG sites. Importantly, previous studies reported no substantial difference in PA between twins and singletons (7,59). As the EWAS was performed on blood samples several years after puberty was completed, the main limitation of this study is that it is unclear whether the methylation profile associated with PDS or PA was formed before, during, or after puberty. Although not revealing clear causal characteristics, the methylation profile in the peripheral blood of young adults is associated with pubertal timing and development, making the associated genes, enriched pathways and linked diseases potential intriguing subjects for future investigation. Furthermore, incorrect reporting of pubertal age, especially in males, could be a source of noise and inconsistency within the analyses (60). In addition, while the self-reported PDS is not the gold standard for measuring pubertal development, it is considered reliable (61). Furthermore, clinical assessments, such as Tanner staging (62,63), are too costly for study designs and sample sizes such as ours. Another limitation was the use of two different methylation profiling platforms, reflecting the technological development of array-based methylation analysis. This resulted in high heterogeneity of some of the CpG sites associated with PDS, denoting perhaps a less reliable association. Nevertheless, the overall heterogeneity between the 450K and EPIC platforms was very low.

## CONCLUSIONS

By identifying CpG sites associated with PDS and PA, we found potential genes, pathways and diseases relevant to puberty in an epigenetic context and complemented the previously reported genetic characterisation of pubertal timing. Our comprehensive analyses on puberty-related CpGs, such as twin modelling, assessment of environmentally driven associations or genetics underlying the association, highlight the role of genetics and unique environment in pubertal timing and development. In addition, we provide evidence for DNA methylation being a putative mechanistic link between genotypes and puberty, as well as puberty-related diseases. Furthermore, our within-pair effects are consistent with a causal environmentally mediated relationship between pubertal development and methylation. To further advance our understanding, it will be essential to utilise longitudinal data from multiple cohorts, such as the Adolescent Brain Cognitive Development Study. By doing so, we could acquire additional insights into the chronological methylation patterns of CpGs associated with PDS and PA, enabling us to deepen our knowledge of the intricate mechanisms underlying puberty and its implications for health and disease.

## DECLARATIONS

### Ethics approval and consent to participate

The Finnish Twin Cohorts’ data collection and analysis were approved by the Ethics Committee of the Helsinki University Central Hospital (Dnro 249/E5/01, 270/13/03/01/2008, 154/13/03/00/2011). Written informed consent was provided by the participants prior to data collection.

### Consent for publication

Not applicable

### Availability of data and materials

The Finnish Twin Cohort datasets used in the current study are located in the Biobank of the Finnish Institute for Health and Welfare, Helsinki, Finland. All biobanked data are publicly available for use by qualified researchers following a standardised application procedure (https://thl.fi/en/web/thl-biobank/for-researchers).

## Authors’ contributions

MO, JK and ES conceptualised the study. ES performed the data analysis, was involved in the visualisation of results, and wrote the first draft of the manuscript. SMZ performed the twin analyses and participated in writing. MKY was involved in data analysis and visualisation. AH performed data preprocessing and normalisation and contributed to data analysis. MO and JK acquired the data, supervised the work and participated in writing. All authors critically revised the manuscript for important intellectual content and read and approved the final manuscript.

## Acknowledgements

We express our gratitude to Teemu Palviainen for his assistance with data management and curation and to Mikaela Hukkanen for her advice and discussions on IPA. We also thank Dr. Giovanna Chiorino for her invaluable advice and insightful discussions. Finally, we thank the FIMM Technology Centre supported by HiLIFE and Biocenter Finland for DNA methylation data generation services, as well as all FinnTwin12 and FinnTwin16 participants and their families included in this study.

## Additional Files

**Additional File 1.docx** - Supplementary methods on EWAS designs and twin modelling.

**Additional File 2.pdf** - **Supplementary Figure S1.** Manhattan plots on p values of meta-analysed EWAS models on 450K and EPIC platforms on PDS at age 12 A) in males and B) females, C) combined, on PDS at age 14 in D) males and females combined, and on PA in E) males and F) females. **Supplementary Figure S2.** QQ plots on p values of meta-analysed EWAS models on 450K and EPIC platforms on PDS at age 12 A) in males and B) females, C) combined, on PDS at age 14 in D) males and females combined, and on PA in E) males and F) females.

**Additional File 3.xlsx** - **Table S1.** Sample sizes and final numbers of complete twin pairs included in 450K and EPIC platforms for EWAS designs on PDS and PA. **Table S2.** Details on univariate twin modelling results on the 14 CpG sites. The estimates for total phenotypic variance and mean, as well as the coefficients of the covariates (age and sex), are shown. **Table S3.** Individual associations and within-twin pair methylation differences of the 58 CpG sites identified in the EWAS. **Table S4.** Results of qualitative and quantitative sex differences for omnibus tests and specific parameters on CpG sites. **Table S5.** CpG sites associated with pubertal development scale (PDS) with a standardised effect size > |0.13| stratified by the model. **Table S6.** CpG sites associated with pubertal age (PA) in males or females with a standardised effect size > |0.13|. **Table S7.** CpG sites associated with pubertal development scale (PDS) or pubertal age (PA) with a standardised effect size > |0.13| enriched in canonical pathways from the Ingenuity Pathway Analysis. **Table S8.** CpG sites associated with pubertal development scale (PDS) or pubertal age (PA) with a standardised effect size > |0.13| enriched in diseases and functions from the Ingenuity Pathway Analysis. **Table S9.** IPA results on meQTLs (p < 5 x 10-8) of CpG sites associated with pubertal development scale (PDS) or pubertal age (PA). Both the Ingenuity Canonical pathways and diseases/functions annotations are included.

