## Additional file 1 for "DNA methylation sites in early adulthood characterised by pubertal timing and development: A twin study"

**Supplementary methods on EWAS designs and twin modelling**

**EWAS designs**

Due to BMI and PDS/PA being functionally intertwined, adding it as a covariate would inherently remove a significant portion of the true association between PDS/PA and methylation values. Therefore, BMI was not included as a covariate in any of the models. Model designs and included covariates are found in Table M1.

**Table M1**. List of EWAS models used in the study. The designs were the same for the EPIC and 450K runs. Meta-analysis was performed on the output of each EWAS design on 450K and EPIC to obtain the merged standardized effect sizes and p-values

| **Model name** | **Covariates** | **Sex** | **Cohort** |
| --- | --- | --- | --- |
| PDS 12 – MF | PDS_12 + age at subproject + date + row + CD8T + CD4T + Bcell + Mono + NK + Neu + smoking + alcohol consumption + sex | M + F | FT12_12 |
| PDS 12 – M | PDS_12 + age at subproject + date + row + CD8T + CD4T + Bcell + Mono + NK + Neu + smoking + alcohol consumption | M | FT12_12 |
| PDS 12 – F | PDS_12 + age at subproject + date + row + CD8T + CD4T + Bcell + Mono + NK + Neu + smoking + alcohol consumption | F | FT12_12 |
| PDS 14 – MF | PDS_14 + age at subproject + date + row + CD8T + CD4T + Bcell + Mono + NK + Neu + smoking + alcohol consumption + sex | M + F | FT12_14 |
| PDS 14 – M | PDS_14 + age at subproject + date + row + CD8T + CD4T + Bcell + Mono + NK + Neu + smoking + alcohol consumption | M | FT12_14 |
| PDS 14 – F | PDS_14 + age at subproject + date + row + CD8T + CD4T + Bcell + Mono + NK + Neu + smoking + alcohol consumption | F | FT12_14 |
| PA – M | PA_16_17 + age at subproject + date + row + CD8T + CD4T + Bcell + Mono + NK + Neu + cohort + smoking + alcohol consumption | M | FT12_17/ FT16_16 |
| PA – F | PA_16_17 + age at subproject + date + row + CD8T + CD4T + Bcell + Mono + NK + Neu + cohort + smoking + alcohol consumption | F | FT12_17/ FT16_16 |

**Twin Modelling**

The first step in twin modelling is to test the mathematical assumptions underlying the univariate model; this is done by fitting a saturated model and comparing it to various reduced models via a likelihood ratio test. The primary assumptions here are 1) the phenotype mean does not differ across twin order, 2) the phenotype variance does not differ across twin order, and 3) the phenotype mean and variance do not differ across zygosity. The assumptions would be considered met if the restricted model was not significantly different from the saturated model as indexed by a likelihood ratio test. CpG meeting these three assumptions were included in further twin analyses.

The next step in twin modelling is to determine the most appropriate variance components underlying each phenotype. Trait variance is composed of additive genetic effects (A), non-additive genetic effects (epistasis for example, D), familial or shared environmental effects that act to make twins more similar to each other (C), and unique environmental effects that act to make twins differ from each other (E). The E term also absorbs measurement error; therefore, it is not removed from models. Under a classical twin design including monozygotic (MZ) and dizygotic (DZ) twins, only an ACE or ADE model can be identified, not a full ACDE model. This ACDE model would require an extended family design to estimate all four variance components simultaneously. Thus, studies of MZ and DZ twins can separately estimate ACE and ADE models and compare model fit to determine which components best explain the phenotypic variance. Reduced models (ex., AE, CE) can be fit by fixing one component to 0 and comparing the reduced model to the full model containing that component freely estimated. As detailed in the Supplemental Results, here we compared all possible models: ACE, ADE, AE, CE, DE, and E only models.

.
