## Additional file 2 for "DNA methylation sites in early adulthood characterised by pubertal timing and development: A twin study"

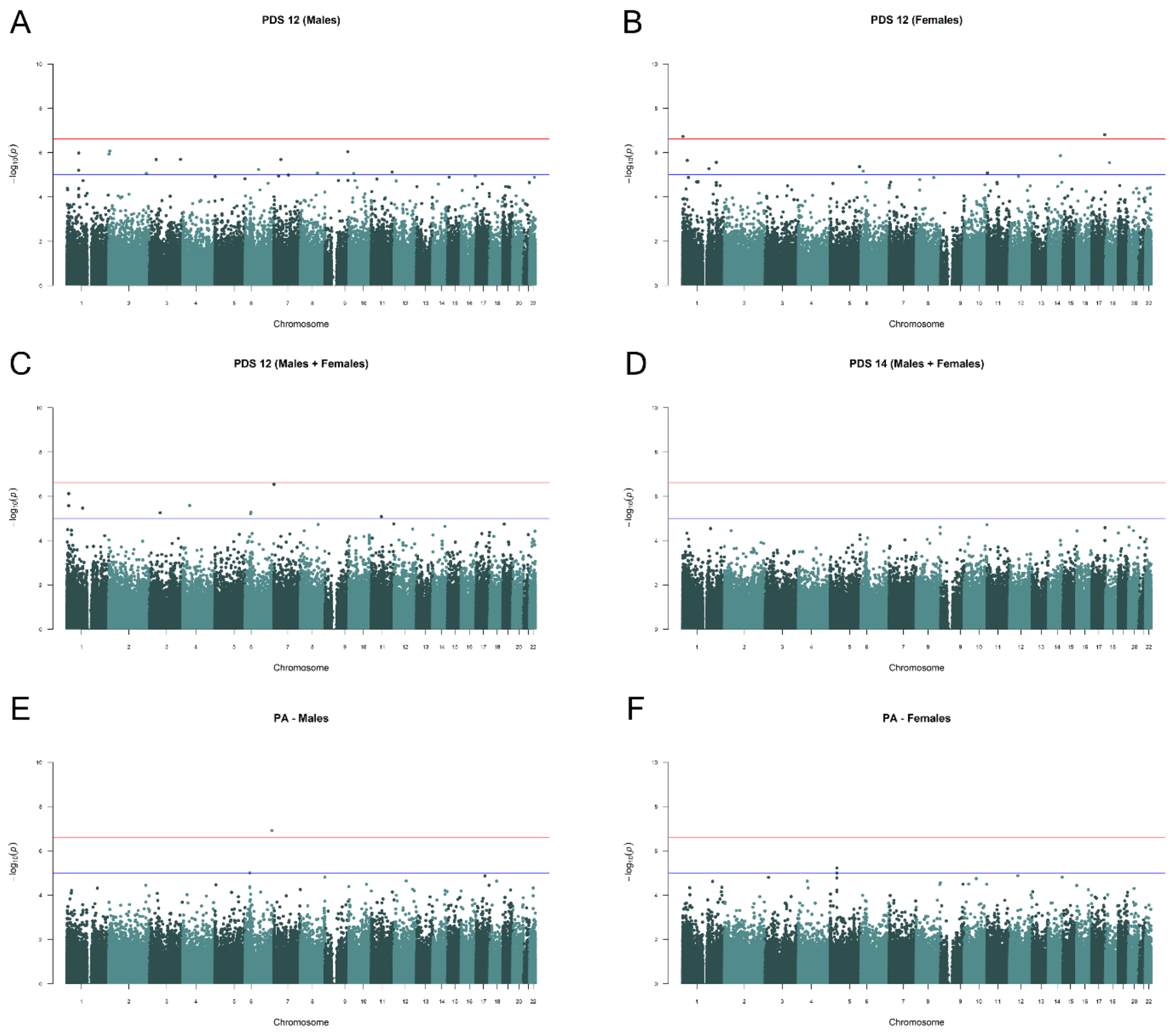


**Supplementary Figure S1.** Manhattan plots on p-values of meta-analysed EWAS models on 450K and EPIC platforms on PDS at age 12 A) in males and B) females, C) combined, on PDS at age 14 in D) males and females combined, and on PA in E) males and F) females

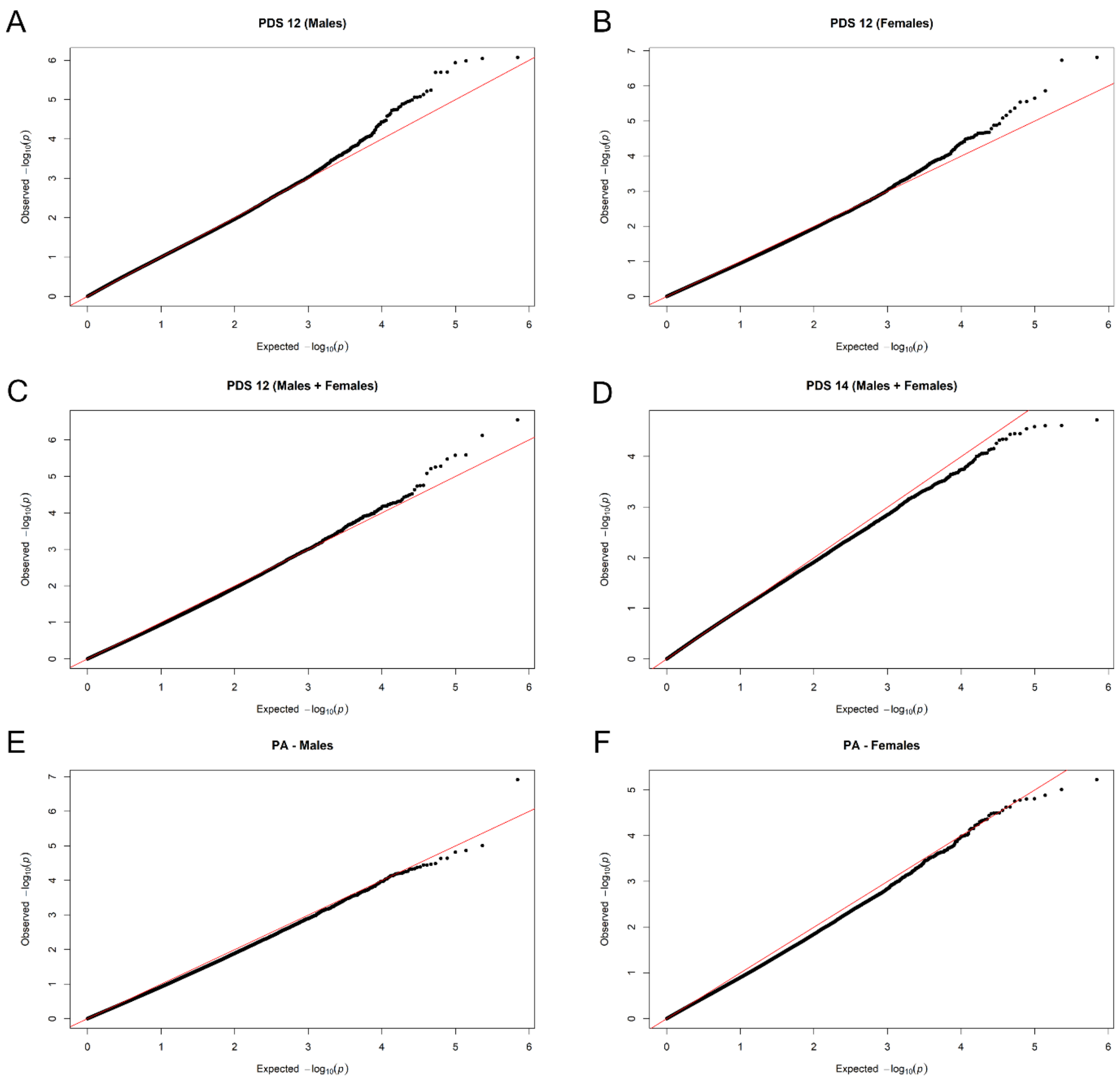


**Supplementary Figure S2.** QQ plots on p-values of meta-analysed EWAS models on 450K and EPIC platforms on PDS at age 12 A) in males and B) females, C) combined, on PDS at age 14 in D) males and females combined, and on PA in E) males and F) females
